# Increased glucose availability sensitizes pancreatic cancer to chemotherapy

**DOI:** 10.1101/2022.04.29.490090

**Authors:** Ali Vaziri-Gohar, Jonathan J. Hue, Hallie G. Graor, Ata Abbas, Mehrdad Zarei, Omid Hajihassani, George Titomihelakis, John Feczko, Moeez Rathore, Rui Wang, Mahsa Zarei, Maryam Goudarzi, Renliang Zhang, Belinda Willard, Li Zhang, Gi-Ming Wang, Curtis Tatsuoka, Joseph M. Salvino, Ilya Bederman, Henri Brunengraber, Costas A. Lyssiotis, Jonathan R. Brody, Jordan M. Winter

## Abstract

Pancreatic cancer (Pancreatic Ductal Adenocarcinoma; PDAC) is highly resistant to chemotherapy. Effective alternative therapies have yet to emerge, leaving chemotherapy as the best available systematic treatment. The discovery of safe and available adjuncts that improve chemotherapeutic efficacy would potentially improve survival outcomes. We show that a hyperglycemic state enhances the efficacy of conventional single- and multi-agent chemotherapies against PDAC. Molecular analyses of tumors exposed to relatively high glucose levels revealed that a key metabolic pathway, glutathione biosynthesis, is diminished and underlies chemo-sensitization by enhancing oxidative injury to cancer cells. Inhibition of this pathway under normal conditions phenocopied a hyperglycemic state by enhancing chemotherapeutic efficacy in mouse PDAC, while rescuing the pathway under high glucose abrogated the anti-tumor effects observed with chemotherapy.

---

Although approved multi-agent chemotherapy regimens offer an incremental advance over single-agent treatments, the best available treatment cocktails are only marginally effective, and virtually all PDAC eventually becomes resistant to these treatments (*1, 2*). In patients with metastatic disease, FOLFIRINOX (FFX; folinic acid, 5FU (5-fluorouracil), irinotecan, and oxaliplatin) or gemcitabine plus albumin-bound (nab) paclitaxel confer a survival benefit of four months or less, compared to single-agent gemcitabine (*3, 4*). Consequently, the median survival of patients with metastatic pancreatic cancer in the modern era is 8-11 months, and the five-year survival rate is around 3% (*5*). The relative ineffectiveness of standard chemotherapy (standard of care; SOC) in murine PDAC models mirrors poor outcomes in patients (*6, 7*).

Nevertheless, chemotherapy continues to be the main treatment option against PDAC due to the absence of effective alternative approaches. In light of this treatment void, strategies to increase the potency of available agents have been an intense focus (*8*). Along these lines, a consortium of leading PDAC foundations in North America is co-sponsoring a phase II randomized trial (PASS-01) to identify predictive markers that select patients for the optimal chemotherapeutic regimen (NCT04469556). The identification of readily available, safe, and effective adjuncts that actually enhance the activity of existing chemotherapies would further improve patient survival and avoid the financial, technical, and regulatory challenges associated with traditional drug development approaches.

One possible reason for poor anti-tumor activity with chemotherapy is that pancreatic tumors are extremely desmoplastic (*9*). Beyond drug delivery considerations (*6, 10*), this pathologic feature is associated with low microvascular density (*6, 11*), tissue hypoxia, and steep nutrient gradients (*12, 13*). To thrive under such harsh conditions, cancer cells require specific molecular adaptations, including enhanced utilization of alternative energy substrates (*10, 14, 15*), optimized mitochondrial function (*13, 16*), and improved handling of reactive oxygen species (*13*). Our group previously demonstrated that an RNA-binding protein, HuR, translocates to the cytoplasm under acute metabolic stress to support these and other survival pathways (*17*). We showed further that these adaptive mechanisms lead to acquired chemoresistance. Due to the acute stress response and associated HuR survival network, chemotherapy is less effective under nutrient limitation in cell culture and mouse PDAC models. Consistent with these observations, patients receiving chemotherapy experienced worse survival after surgery when their peripheral glucose levels were in the normal range (i.e., leading to nutrient-deprived and more resistant tumors), as compared to hyperglycemic patients (harboring tumors exposed to relative glucose abundance) (*17*). These findings may indicate that serum glucose levels predict chemotherapy response; for example, higher glucose predicts chemo-sensitivity. A more intriguing take-away from this research is that glycemic status is modifiable to potentially improve PDAC response to chemotherapy. Thus, we hypothesized that forced hyperglycemia sensitizes PDAC to chemotherapy and further aimed to understand the molecular underpinnings of this phenomenon.

We first validated the prior and aforementioned retrospective clinical study (*17*) performed on a cohort of patients with localized PDAC. In the present series, we examined the impact of glycemic status on patients with metastatic PDAC. Approximately 33% of patients presented with diabetes mellitus, consistent with historical experience (*18*). There were no appreciated demographic differences between normal and high glucose patients (**Extended Data Table 1**). A greater proportion of patients in the high glucose group already carried a documented clinical diagnosis of diabetes mellitus at the time of PDAC diagnosis, as compared to the normal glucose group (56.2% vs. 14.8%, *P*<0.001). Median pre-diagnosis (137 vs. 105 mg/dL, *P*<0.001) and treatment-period glucose levels (158 vs. 109 mg/dL, *P*<0.001) were higher among patients in the high glucose group. The median CA 19-9 level, a prognostic marker commonly used to reflect disease burden and PDAC aggressiveness, was similar between groups at diagnosis, although it trended towards a higher value in the high glucose group (2439.5 U/mL high glucose vs 1294.5 U/mL normal glucose, *P*=0.224). First-line chemotherapy regimens were similar as well, with the majority of patients in the entire cohort receiving multi-agent standard-of-care regimens (*P*=0.881). The median overall survival among all patients who completed at least two cycles of chemotherapy was approximately 9.8 months in the overall cohort (IQR: 6.3, 14.9 months), which is on par with historical clinical trial data (*3, 4*). Multivariable Cox proportional hazards regression demonstrated that patients in the high glucose levels had an associated survival benefit relative to those in the normal glucose group (HR=0.61, 95% CI 0.41- 0.92, *P*=0.02) (**Fig. 1a**), after controlling for medical comorbidities, performance status, site of metastatic disease, CA 19-9 value at diagnosis, total number of chemotherapy cycles, and chemotherapy regimen (**Extended Data Table 2**). Notably, no associated survival difference was observed based on glucose levels in an independent cohort of metastatic patients who did not receive treatment (HR=0.99, 95% CI 0.64-1.53, *P*=0.97) (**Fig. 1b**), suggesting that the interaction with glycemic status may be present only for patients who received chemotherapy. Paired with prior previously published cell culture data by our group (*17*), these clinical data indicate that a high-glucose state sensitizes PDAC to chemotherapy.

**Figure 1.**
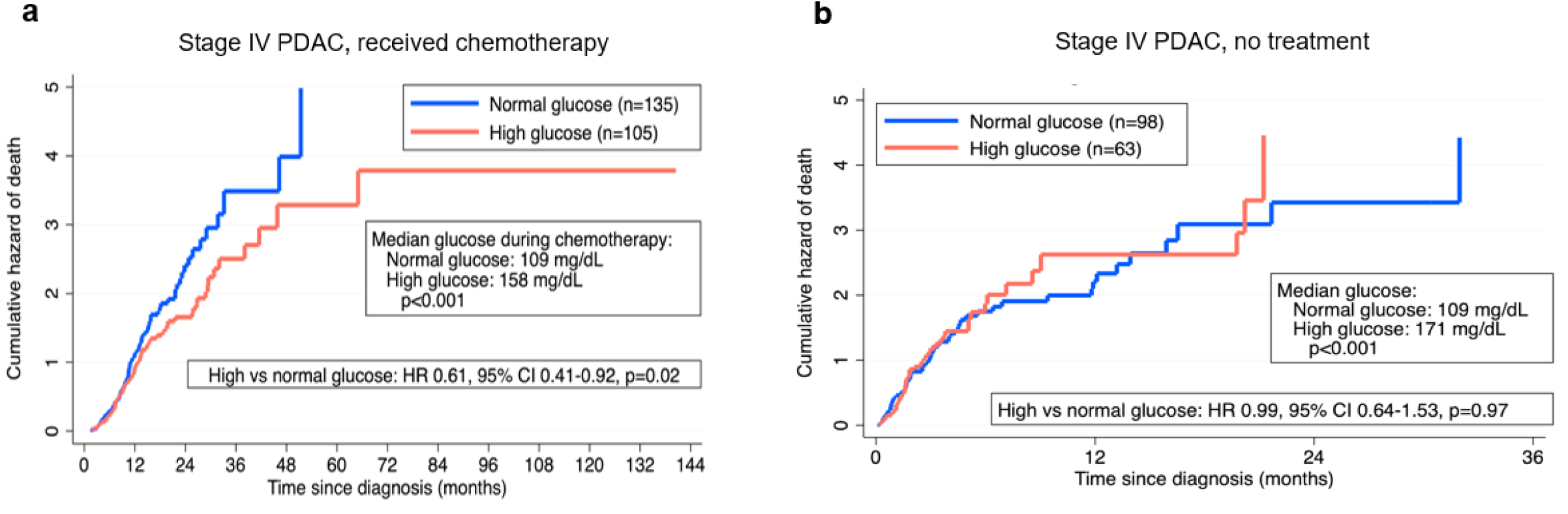
Enhanced chemotherapy response with hyperglycemia in patients with stage IV PDAC. Nelson-Aalen cumulative hazard curves of patients with metastatic PDAC, stratified by glycemic status among patients who received ≥ 2 cycles of chemotherapy (**a**) and those who did not receive any treatment (**b**). Displayed mortality curves reflect multivariable Cox proportional hazards regression (Extended Data Table 2).

We followed up on these findings with a series of controlled studies in mouse PDAC models. First, we induced hyperglycemia pharmacologically using streptozotocin (STZ). This drug chemically ablates β-islet cells in the pancreas to induce pancreatogenic diabetes. Since this model results in extremely elevated and debilitating glucose levels, titration to non-toxic levels of hyperglycemia was achieved with daily injections of long-acting insulin glargine (**Fig. 2a**). Hyperglycemia was also induced through dietary changes in independent experiments. Specifically, mice were allowed to consume 30% dextrose water (D30) *ad libitum*. This hyperglycemic model was previously validated by our lab through serial peripheral glucose monitoring, along with intra-tumoral glucose measurements (*13*). PDAC xenografts in hyperglycemic nude mice reproducibly exhibited greater sensitivity to single-agent chemotherapy (**Fig. 2b**). Peripheral glucose levels and mouse weights associated with each hyperglycemic model are provided (**Fig. 2c, d)**, and demonstrate increased peripheral glucose levels with stable body weights, echoing previous reports (*13*). Consistent with the above results from MiaPaCa-2 xenografts, patient-derived xenografts were also more responsive to chemotherapy in hyperglycemic mice (**Fig. 2e**). As observed previously with patients in the absence of chemotherapy (**Fig. 1b**), no differences in growth rates were observed in vehicle-treated mice (**Fig. 2b, e**).

**Figure 2.**
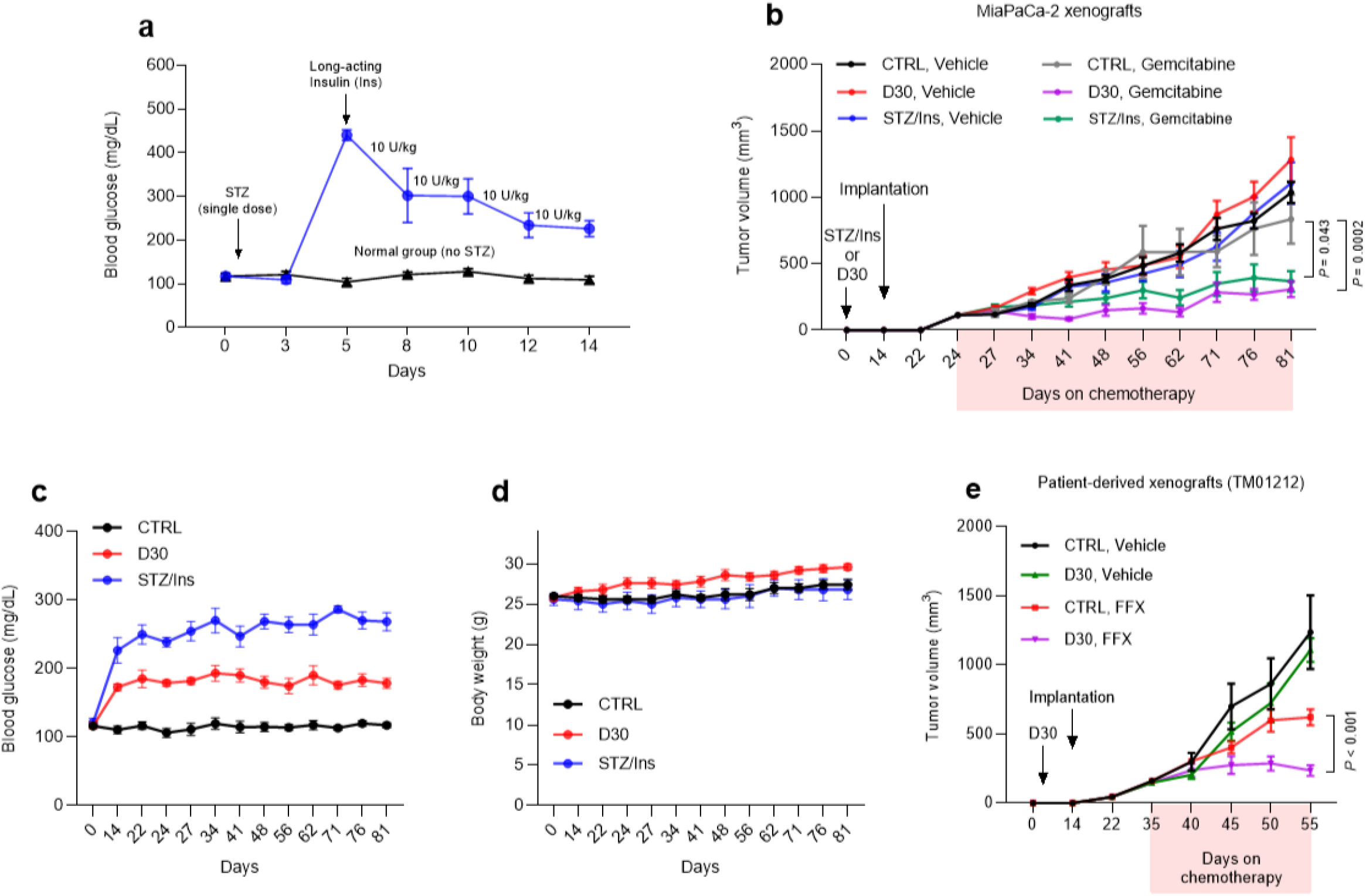
Hyperglycemia mice respond better to conventional chemotherapy. **a**. Pharmacologically induced hyperglycemia in nude mice with a single dose of streptozotocin (STZ, 120 mg/kg) followed by daily subcutaneous insulin glargine injections to keep glucose levels in a non-toxic range. **b**, Xenograft growth of MiaPaCa-2 cells treated with gemcitabine (n=5 per group). **c, d**, Peripheral glucose levels (**c**) and body weights (**d**) of nude mice that received normal water, D30 or STZ with insulin glargine (STZ/Ins). Each dot represents the mean of weekly measurements of blood glucose per group. **e**, Patient derived xenograft (TM01212) treated with FOLFIRINOX (n=5 per group). Data are provided as mean ± s.e.m. Longitudinal mixed models were fit for tumor size growth, and time by treatment interactions were assessed (**b**,**e**).

Providing D30 water to C57BL/6J mice similarly increased peripheral blood glucose and intra-tumoral glucose levels relative to normoglycemic mice (**Extended Data Fig. 1a-c**). Consistent with prior study (*19*), these results indicating the intratumoral nutrient levels can be influence by surrounding conditions. Metabolite studies of tumors revealed a substantial increase in fatty acids in the hyperglycemic group (**Extended Data Fig. 1c**). On the other hand, D30 consumption resulted in a significant reduction in the abundance of TCA (tricarboxylic acid) cycle-associated metabolites in tumors (**Extended Data Fig. 1c**). In these tumors, principal component analysis (PCA) also revealed a global change in transcriptomes with hyperglycemia (**Extended Data Fig. 1d**), reflected by 1843 differentially expressed genes (**Extended Data Fig. 1e**). Among other pathways, transcriptomic analyses demonstrated that glycolysis- and fatty acid synthesis-related pathways were induced in tumors of mice in the D30 group, paralleling the metabolite studies (**Extended Data Fig. 1f, g**). In total, results from two separate mass spectrometric analyses, GC-MS and LC-MS, demonstrated that fatty acids accumulated in tumors exposed to a hyperglycemic state (**Extended Data Fig. 1h** and **1c** respectively). Activation of the synthetic macromolecular machinery did not however translate into increased tumor growth or cancer cell proliferation in the absence of chemotherapy. Similarly, gene set enrichment analyses did not reveal a change in mitotic spindle assembly or DNA replication in high-glucose tumors (**Extended Data Fig. 1i**).

More granular assessment of transcriptome changes indicated a general induction of oxidative stress-associated genes under high glucose conditions. While this observation likely reflects an adaptive response by cancer cells to protect against free radicals, a number of redox genes were significantly downregulated under these conditions (**Fig. 3a**). Notably, the catalytic subunit of glutamate-cysteine ligase (GCLC) was considerably reduced in KPC tumors under hyperglycemia (**Fig. 3b-d**). This enzyme is the first and rate-limiting enzyme for *de novo* glutathione biosynthesis and enzymatically catalyzes the union of glutamate and cysteine. Along these lines, GSH/GSSG levels were reduced in D30-treated tumors in mice (**Fig. 3e**). Similar findings were observed with human PDAC cells and xenografts under high glucose conditions (**Fig. 3f, g**). A reduction in the glutathione precursors, glutamine and glycine, further reveals dysregulation of this pathway in tumors under high glucose abundance (**Extended Data Fig. 1c**). Taken together, these data indicate that GCLC expression and reduced glutathione levels are diminished in niches with higher relative glucose abundance (*12, 13*).

**Figure 3.**
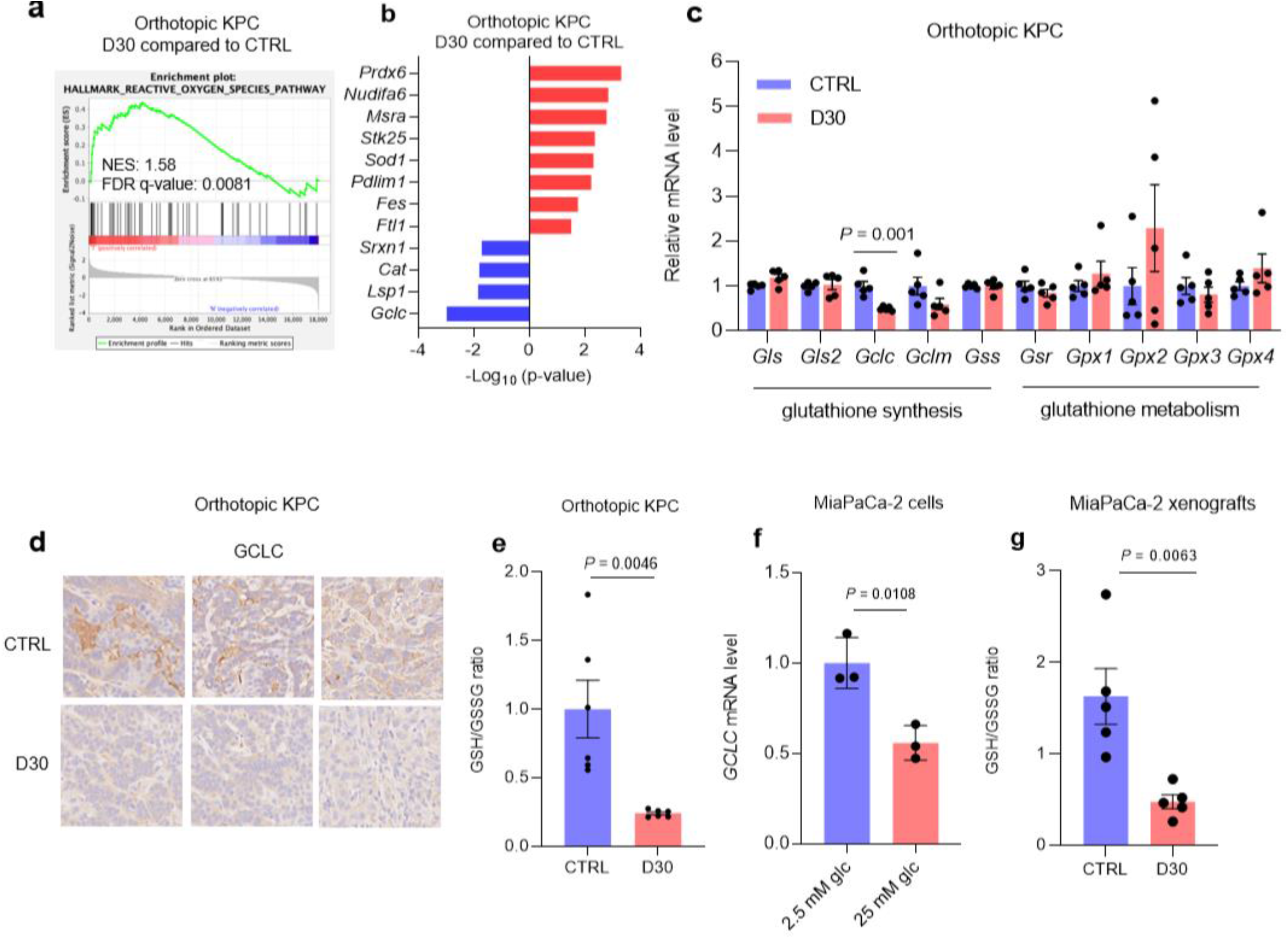
Reduced *de novo* glutathione synthesis in pancreatic cancer exposed to a relative high glucose state. **a-c**, GSEA of differentially expressed genes associated with oxidative stress (**a**), significantly altered genes associated with redox metabolism (**b**) and qPCR analysis of glutathione-associated enzymes (**c**) in KPC orthotopic tumors under the indicated conditions (n=5 orthotopic tumors). **d**, qPCR analysis of *GCLC* transcripts in MiaPaCa-2 cells under the indicated conditions for 48 hours (n=3 independent experiments). **e**, Immunolabeling of GCLC in independent KPC orthotopic tumors receiving D30 water versus control. **f, g**, GSH/GSSG ratio in KPC orthotopic tumors (**f**) and MiaPaCa-2 xenografts (**g**) under the indicated conditions (n=5 per group). Data are provided as mean ± s.e.m (**c, e, g**).

The antioxidant and chemotherapy-resistant properties of GSH are well-described (*20-22*). Thus, reduced levels in tumors exposed to a high-glucose state may underlie improved chemotherapy response in this context. As proof of concept, chemotherapeutic agents induced substantial ROS levels under high glucose, but not under low glucose conditions (**Fig. 4a**). siRNA against GCLC abrogated the chemotherapy resistance observed in parental or siCTRL-transfected PDAC cells cultured under low glucose conditions (**Fig. 4b**).

**Figure 4.**
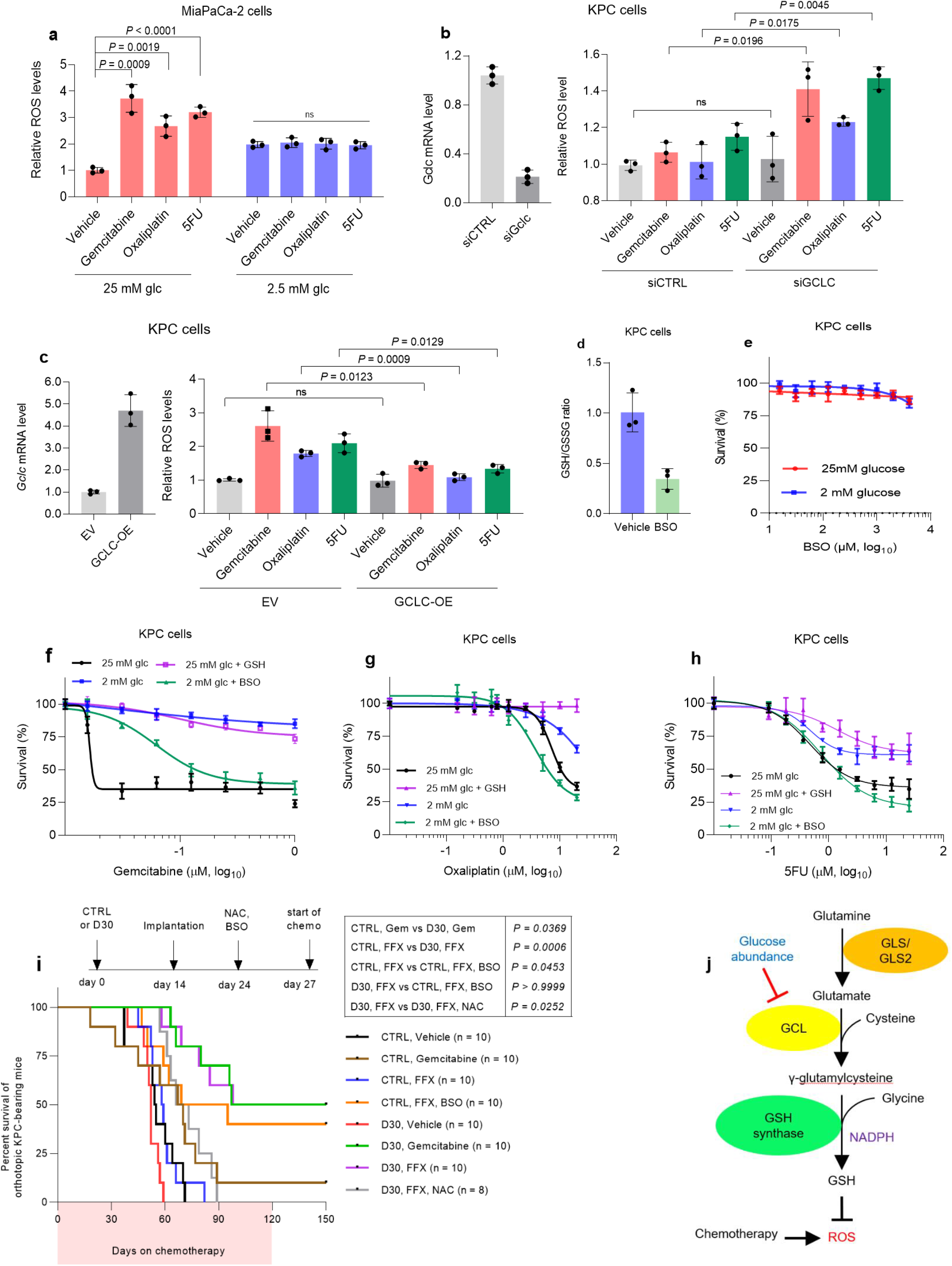
Downregulation or inhibition of glutathione synthesis enhances chemotherapeutic efficacy in PDAC. **a**, Relative ROS levels in MiaPaCa-2 cells cultured in low (2.5 mM) or high (25 mM) glucose for 30 hours, followed by chemotherapy administration (Gemcitabine (100 nM), Oxaliplatin (1 µM), 5FU (1 µM)) for an additional 16-18 hours (n= 3 independent experiments). Relative ROS levels in murine KPC cells transiently transfected with siRNAs (control non-targeting or against *GCLC*) cultured in low glucose (2 mM) (**b**) or transiently transfected with *GCLC* overexpressing plasmid cultured in high glucose (25 mM) (**c**) conditions for 30 hours followed by chemotherapy administration (Gemcitabine (100 nM), Oxaliplatin (1 µM), 5FU (1 µM)) for an additional 16-18 hours (n= 3 independent experiments). **d**, GSH/GSSG ratio in MiaPaCa-2 cells treated with vehicle or BSO (0.5 mM) under low glucose (2 mM) condition for 48 hours (n=3 independent experiments). **e**-**h**, Relative survival of KPC cells at the indicated conditions for five days. For experiments under low glucose conditions, cells were cultured in media containing 2 mM glucose for 30 hours prior to chemotherapy treatment. GSH (reduced glutathione, 4 mM) was co-administered with chemotherapy. BSO (L-Buthionine sulfoximine, 0.5 mM) was co-administered with low glucose medium (n= 3 independent experiments). **i**, Survival of C57BL/6J mice bearing orthotopic murine KPC pancreatic tumors with the indicated treatments. **j**, Depiction of the *de novo* glutathione synthesis pathway and the effects of relative glucose abundance. Data are provided as mean ± s.d. (**a**-**h**). Survival distributions were estimated using Kaplan-Meier estimation and compared by log-rank tests, and adjusted p-values are reported using Holm’s method for multiple comparisons (**i**).

Conversely, GCLC overexpression minimized ROS induction under high glucose in cultured PDAC cells treated with conventional chemotherapy (**Fig. 4c**). A GCLC inhibitor, BSO (L-buthionine sulfoximine) (*23*), reduced GSH/GSSG levels in cultured PDAC cells under low glucose, as expected (**Fig. 4d**). Despite the adverse effect on reductive power, the drug had a negligible effect on cancer cell survival in mice as a monotherapy (**Fig. 4e**), in keeping with a prior report (*24*). However, paired with an ROS-inducing agent (i.e., chemotherapy), BSO neutralized the chemotherapy-resistance effect of low glucose conditions on PDAC cells. BSO-associated cytotoxicity under low glucose even trended towards levels observed with chemotherapy under high glucose conditions (**Fig. 4f-h**). For these experiments, supplemental reduced glutathione (GSH) rescued PDAC cells treated with chemotherapy in the high-glucose state (**Extended Data Fig. 4f-h**).

Standard-of-care, FOLFIRINOX, had no anti-tumor effect in a syngeneic orthotopic PDAC model at the indicated dosing schedule (**Fig. 4i**), and as previously observed (*7*). The addition of BSO to chemotherapy improved survival (median survival: 58 vs. 69 days, *P*=0.0453 **(Fig. 4i**)). Still, anti-PDAC activity was greatest with FOLFIRNOX treatment in hyperglycemic mice (median survival: 58 vs. 98, *P*=0.0006 (**Fig. 4i**). The antioxidant and glutathione precursor, NAC, abrogated the sensitizing benefit effect of hyperglycemia to chemotherapy, underscoring the role of oxidative injury to tumors (**Fig. 4i**). Similar results were observed with gemcitabine, where a hyperglycemic state sensitized orthotopic tumors to treatment (median survival: 67 vs. 97 days, *P*=0.0369 (**Fig. 4i**)).

The *GCLC* mRNA transcript is a known regulatory target of the acute stress response protein, HuR (*25*). No significant change was observed with HuR expression in orthotopic PDAC tumors (**Extended Data Fig. 2a**). Therefore, we hypothesized that *GCLC* downregulation under high glucose conditions was a function of reduced HuR subcellular localization to the cytoplasm, as previously observed in cell culture models under these conditions (*17, 26*). Indeed, cytosolic HuR immunolabeling was reduced in tumors exposed to a high glucose state compared to tumors in normoglycemic mice. In contrast, nuclear labeling was especially prominent in D30 treated tumors (**Extended Data Fig. 2b, c**). These findings suggest that HuR activation and subcellular localization to the cytoplasm leads to target transcript destabilization, and reduced *GCLC* expression (*25*), as reported by others and validated in cultured PDAC cells (**Extended Data Fig. 2d**).

Based on these reported findings, we offer a model where reduced GCLC expression under relative glucose abundance potentially underlies PDAC sensitivity to chemotherapy (**Fig. 4j**). Moreover, these data provide a rationale for two independent, but complementary therapeutic strategies, as illustrated here in pre- clinical experiments. Each is testable in the clinical setting. First, tumors can be theoretically ‘primed’ for chemotherapy through a forced hyperglycemic state, carefully controlled by dextrose administration (e.g., dextrose infusion with rigorous inpatient glucose monitoring) at the time of chemotherapy infusion. Second, the addition of BSO to standard chemotherapy, especially in patients with normal glucose levels, would counteract elevated GCLC levels typically present in PDAC under glucose limiting conditions in tumors. The safety of BSO with chemotherapy has already been established in patents with other tumor types (*27, 28*), and strengthens the rationale to pursue this approach in patients with PDAC. Together, these observations offer an approach to improve chemotherapeutic efficacy and survival outcomes in patients with PDAC.

**Extended Data Table 1.** Demographic and clinical data of patients with stage IV pancreatic ductal adenocarcinoma, who received chemotherapy or supportive care (no chemotherapy), stratified by normal or high glucose.

|  | Received chemotherapy |  |  | Did not receive chemotherapy |  |  |
| --- | --- | --- | --- | --- | --- | --- |
|  | Normal glucose | High glucose | P value | Normal glucose | High glucose | P value |
| N | 135 (56.3%) | 105 (43.7%) | --- | 98 (60.9%) | 63 (39.1%) | --- |
| Age, years (median, IQR) | 67 (58, 72) | 65 (58, 74) | 0.94 | 72 (62, 80) | 73 (65, 81) | 0.66 |
| Male sex | 67 (49.6%) | 54 (51.4%) | 0.78 | 48 (49.0%) | 31 (49.2%) | 0.98 |
| White race | 99 (73.3%) | 75 (71.4%) | 0.90 | 68 (69.4%) | 40 (63.5%) | 0.08 |
| ECOG performance status |  |  | **0.04** |  |  | 0.13 |
| ≤1 | 94 (69.6%) | 76 (72.4%) |  | 23 (23.5%) | 17 (27.0%) |  |
| ≥2 | 13 (9.6%) | 18 (17.1%) |  | 21 (21.4%) | 21 (33.3%) |  |
| Not recorded | 28 (20.7%) | 11 (10.5%) |  | 54 (55.1%) | 25 (39.7%) |  |
| Charlson comorbidity score (median, IQR) | 8 (7, 9) | 9 (8, 10) | **0.01** | 9 (8, 10) | 10 (9, 11) | **0.01** |
| Documented diabetes at diagnosis | 20 (14.8%) | 59 (56.2%) | **<0.001** | 21 (22.3%) | 44 (69.8%) | **<0.001** |
| Initial metastatic disease site |  |  | 0.16 |  |  | 0.35 |
| Liver | 80 (59.3%) | 66 (62.9%) |  | 61 (66.3%) | 26 (55.7%) |  |
| Peritoneal cavity | 15 (11.1%) | 9 (8.6%) |  | 3 (3.3%) | 4 (8.2%) |  |
| Multiple sites | 30 (22.2%) | 15 (14.3%) |  | 19 (20.6%) | 11 (27.9%) |  |
| Other, single site | 10 (7.4%) | 15 (14.3%) |  | 9 (9.8%) | 4 (8.2%) |  |
| CA 19-9 at diagnosis, U/mL (median, IQR) | 1294.5 (162.4, 7446.2) | 2439.5 (246.2, 14318.8) | 0.22 | 1910.8 (95.0, 30099.1) | 1715.0 (220.5, 19982.9) | 0.70 |
| First-line chemotherapy regimen |  |  | 0.88 |  |  |  |
| FOLFIRINOX | 51 (37.8%) | 35 (33.3%) |  |  |  |  |
| Gemcitabine with nab-paclitaxel | 38 (28.2%) | 34 (32.4%) |  |  |  |  |
| Gemcitabine | 15 (11.1%) | 10 (9.5%) |  |  |  |  |
| FOLFOX | 20 (14.8%) | 15 (14.3%) |  |  |  |  |
| Other | 11 (8.1%) | 11 (10.5%) |  |  |  |  |
| Number of cycles, first-line (median, IQR) | 3 (1, 6) | 4 (2, 8) | 0.24 | 0 | 0 | --- |
| Total number of cycles (median, IQR) | 5 (2, 10) | 7 (2, 13) | 0.05 | 0 | 0 | --- |

Abbreviations: IQR, interquartile range; ECOG, Eastern Cooperative Oncology Group; FOLFIRINOX (folinic acid, 5-fluorouracil, oxaliplatin, irinotecan); nab, nanoparticle albumin-bound; FOLFOX, folinic acid, 5-fluorouracil, oxaliplatin. Bold values indicate statistical significance.

**Extended Data Table 2.** Multivariable Cox proportional hazards regression analyzing factors associated with overall survival. Separate regressions were performed for patients who received chemotherapy and those who did not receive chemotherapy.

|  | Received chemotherapy |  |  |  | Did not receive chemotherapy |  |  |  |
| --- | --- | --- | --- | --- | --- | --- | --- | --- |
|  | HR | 95% CI |  | P value | HR | 95% CI |  | P value |
| High glucose vs normal glucose | 0.61 | 0.41 | 0.92 | **0.02** | 0.99 | 0.64 | 1.53 | 0.97 |
| Modified Charlson comorbidity index*: $\geq 10$ vs $\leq 9$ | 0.70 | 0.42 | 1.16 | 0.17 | 0.92 | 0.63 | 1.34 | 0.66 |
| Documented diabetes at diagnosis | 1.41 | 0.93 | 2.15 | 0.11 | 1.00 | 0.64 | 1.57 | 0.99 |
| ECOG performance status |  |  |  |  |  |  |  |  |
| $\leq 1$ | Ref. | --- | --- | --- | Ref. | --- | --- | --- |
| $\geq 2$ | 0.72 | 0.41 | 1.26 | 0.25 | 1.25 | 0.75 | 2.09 | 0.40 |
| Unknown | 1.29 | 0.80 | 2.08 | 0.30 | 1.34 | 0.87 | 2.06 | 0.19 |
| Initial metastatic disease site |  |  |  |  |  |  |  |  |
| Liver | Ref. | --- | --- | --- | Ref. | --- | --- | --- |
| Peritoneum | 0.79 | 0.46 | 1.38 | 0.42 | 1.10 | 0.51 | 2.38 | 0.81 |
| Multiple sites | 1.26 | 0.82 | 1.93 | 0.28 | 1.27 | 0.85 | 1.94 | 0.28 |
| Other, single site | 0.48 | 0.28 | 0.85 | **0.01** | 0.62 | 0.32 | 1.18 | 0.15 |
| CA 19-9 at diagnosis |  |  |  |  |  |  |  |  |
| $< 37$ U/mL | 0.98 | 0.64 | 1.51 | 0.94 | 0.92 | 0.56 | 1.51 | 0.73 |
| 37 - 1000 U/mL | 0.76 | 0.50 | 1.14 | 0.18 | 0.59 | 0.35 | 0.98 | **0.04** |
| $> 1000$ U/mL | Ref. | --- | --- | --- | Ref. | --- | --- | --- |
| Unknown | 0.52 | 0.28 | 0.98 | **0.04** | 0.98 | 0.62 | 1.53 | 0.91 |
| Total number of cycles: $\geq 10$ vs $\leq 9$** | 0.15 | 0.10 | 0.22 | **$< 0.001$** | | | | |
| First-line chemotherapy: FOLFIRINOX or gemcitabine with nab-paclitaxel vs other | 0.84 | 0.59 | 1.18 | 0.32 |  |  |  |  |

\*Modified Charlson comorbidity index does not include diabetes mellitus, as this was analyzed separately in multivariable models.
\*\*Patients who received $\leq 1$ cycle of chemotherapy were excluded from analyses.
Abbreviations: IQR, interquartile range; ECOG, Eastern Cooperative Oncology Group; FOLFIRINOX (folinic acid, 5-fluorouracil, oxaliplatin, irinotecan); nab, nanoparticle albumin-bound; HR, hazard ratio; CI, confidence interval; Ref, reference Bold values indicate statistical significance.

**Extended Data Figure 1.**
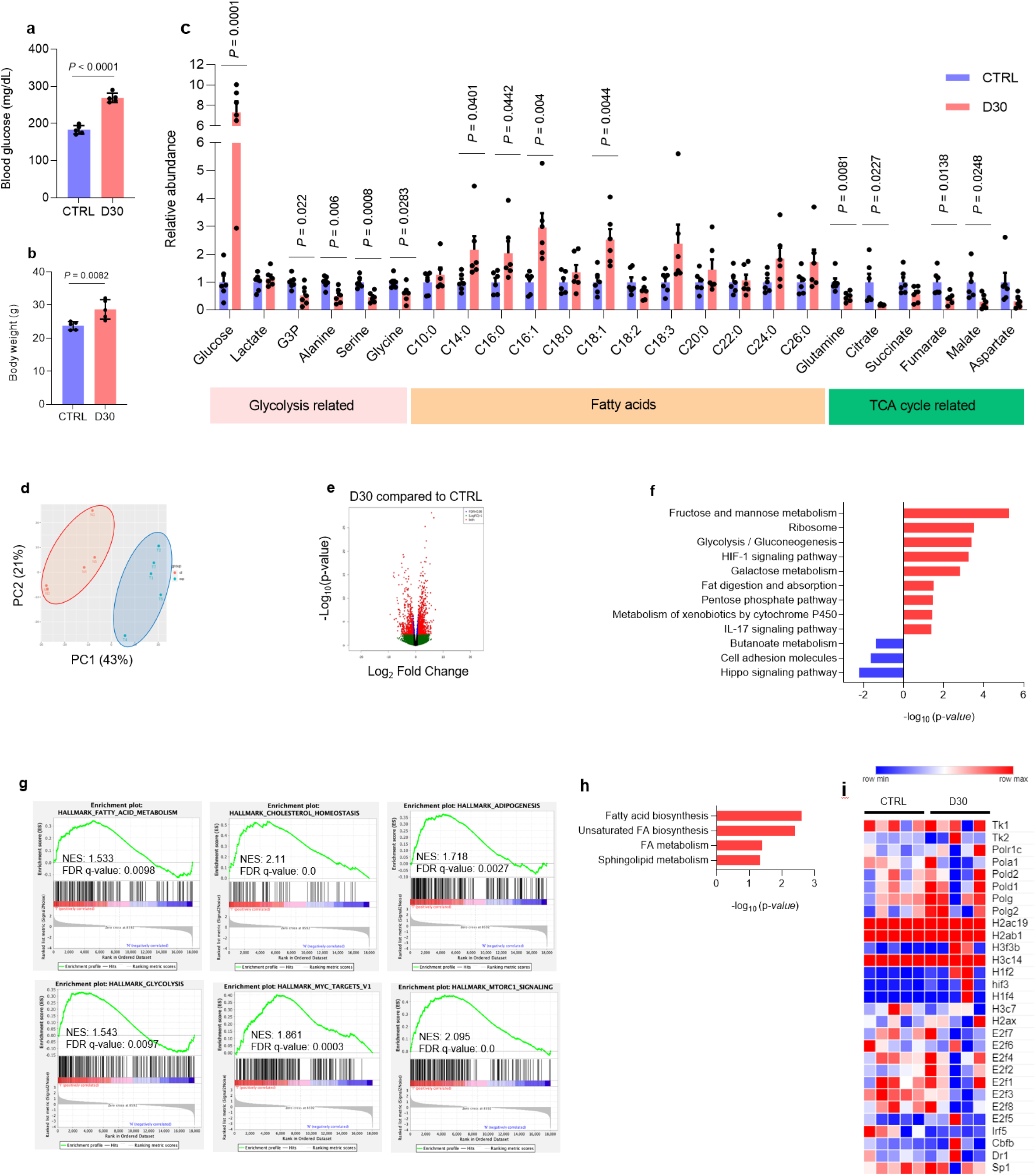
Altered biologic pathways in orthotopic pancreatic cancers from hyperglycemic mice. Peripheral glucose levels (**a**) and body weights (**b**) in C57BL/6J mice receiving D30 water or normal water. **c**, Relative abundance of metabolites in murine KPC orthotopic tumors after D30 water consumption, as compared to tumors in normoglycemic mice consuming regular water for 14 days (n=6 orthotopic tumors). Principle component analysis (**d**) and volcano plot (**e**) in KPC orthotopic tumors under the indicated conditions (n=5 orthotopic tumors). Enriched pathways derived from transcriptomic and metabolomic (**f**), GSEA of differentially expressed genes (**g**) (n=5 orthotopic tumors). **h**, Heatmap of genes associated with DNA replication and cell cycle division in KPC orthotopic tumors under the indicated conditions (n=5 orthotopic tumors). Data are provided as mean ± s.d. (**a, b**) or mean ± s.e.m. (**c**).

**Extended Data Figure 2.**
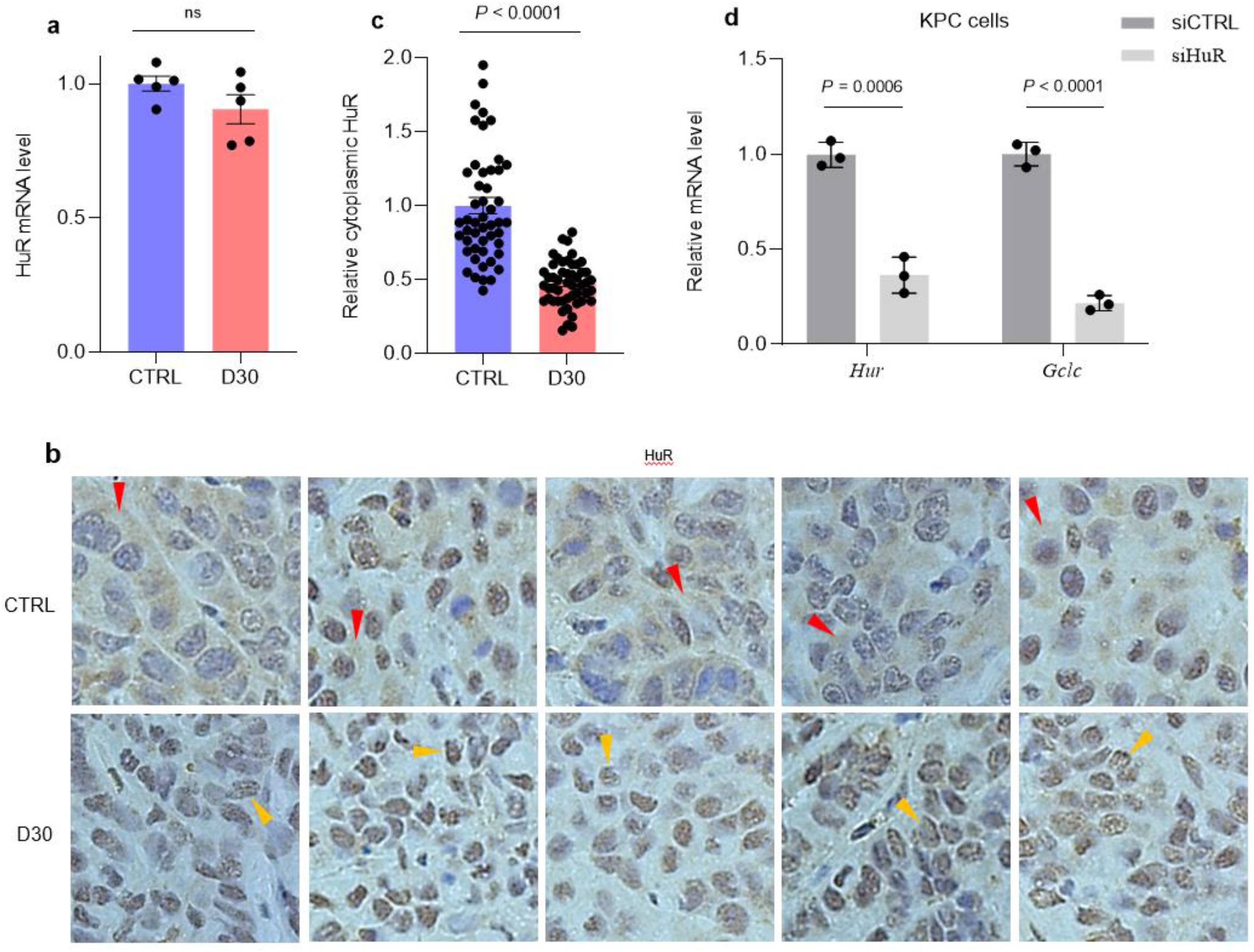
HuR regulates GCLC expression in pancreatic cancer. **a**, qPCR analysis of HuR in KPC orthotopic tumors under the indicated conditions (n=5 tumors per group). Immunohistochemistry immunolabeling of HuR (**b**) and analysis of subcellular localization of HuR (n=50 cells per group) (**c**) in KPC orthotopic tumors from mice receiving D30 water or normal water. Hematoxylin staining was used to stain nuclei. **d**, qPCR analysis of *GCLC* after HuR siRNA silencing in cultured KPC cells (n=3 independent experiments). Data are provided as mean ± s.e.m. (**a, c**) and mean ± s.d. (**d**).

## Methods

### Cell lines, cell culture, and reagents

All cell lines were obtained from ATCC, except murine PC cells (KPC K8484: Kras^G12D/+^; Trp53^R172H/+^; Pdx1-Cre) (*13*). Mycoplasma screening was performed using a MycoAlert detection kit (Lonza). Cell lines were maintained at 37°C and 5% CO2. For standard cell culture, cells were grown in DMEM (25mM glucose, 4mM glutamine), supplemented with 10% FBS, 1% penicillin/streptomycin, and prophylactic doses of Plasmocin (Life Technologies, MPP-01-03). Glucose withdrawal was performed to simulate low glucose conditions characteristic of the PDAC microenvironment. For low-glucose experiments, glucose-free DMEM (Life Technologies, 21013- 024) was titrated to the indicated glucose levels. For rescue experiments, GSH (L-Glutathione reduced, Sigma, G6013), NAC (N-Acetyl-L-cysteine, Sigma, A9165), BSO (L-Buthionine-sulfoximine, Sigma, B2515), and *GCLC* overexpressing plasmid (Origene, MC203908) were used. Chemotherapies included gemcitabine hydrochloride (Sigma, G6423), oxaliplatin (Sigma, O9512), irinotecan (Sigma, I1406), and 5FU (Sigma, F6627).

### Small RNA interference

Oligos were obtained from Life technology siRNA transfections and were performed using Lipofectamine 2000. siRNA gene knockdown validation was determined 72 hours after siRNA transfections via qPCR.

### Cell viability assays

Cell viability was estimated by DNA quantitation using the PicoGreen dsDNA assay (Life Technologies, P7589).

### ROS and GSH/GSSG ratio quantification

Cells were incubated in a 96-well plate with 10µM H2-DCFDA (Life Technology, D399) for 45 minutes in serum-free media to detect total intracellular ROS. GSH/GSSG ratio (Promega, V6611) measurements were performed according to the manufacturer’s instructions. Readouts were normalized to cell number or protein content.

### Quantitative RT-PCR

Total RNA was extracted using PureLink RNA isolation (Life Technologies, 12183025) and treated with DNase (Life Technologies, AM2222). cDNA was synthesized using 1µg of total RNA, oligo-dT and MMLV HP reverse transcriptase (Applied Biosystems, 4387406). All PCR reactions were performed in triplicate; primer sequences are provided in the supplementary section. RT-qPCR acquisition was captured using a Bio-Rad CFX96 and analyzed using Bio-Rad CFX Manager 2.0 software.

### RNA-sequencing and analyses

RNA quality was assessed using an Agilent 2100 Bioanalyzer (Agilent Technologies). Strand-specific RNA-seq libraries were prepared using NEBNext Ultra II Directional RNA Library Prep Kit (NEB, Ipswich, MA) according to the manufacturer’s protocols. RNA-sequencing was performed using 150-bp paired-end format on a NovaSeq 6000 (Illumina) sequencer. FastQC was used to assess RNA-seq quality and TrimGalore was used for adapter and quality trimming. RNA-seq reads were mapped against hg38 using STAR (v2.7.0e) aligner with default parameters. DESeq2 analysis with an adjusted P-value <0.05 was used to derive a list of differentially expressed genes.

### Immunohistochemistry immunolabeling

Samples were preserved in formalin and embedded in paraffin followed by GCLC (Origene, TA507318, Mouse Monoclonal Antibody, Clone ID: OTI1A3, dilution: 1:150) and HuR (Santa Cruz Biotechnology, SC-5261, dilution 1:250) immunolabeling antibodies.

### Metabolic profiling

GC-MS analyses were performed using a Hewlett Packard 5973 Turbo Pump Mass Selective Detector and a Hewlett Packard 6980 Gas Chromatograph equipped with a DB-5ms GC Column (60 m x 0.25 mm x 0.25 um, Agilent Technologies). Samples were weighed and homogenized using Folch method (2:1 chloroform-methanol). For fatty acid measurements, chloroform phase containing TG-bound fatty acids was hydrolyzed using alkaline hydrolysis. Fatty acids were converted to their methyl esters and analyzed by GC-MS. Fatty acids were quantified using a 19:0 fatty acid standard. The methanol/water layer was evaporated to dryness in a Speedvac evaporator at 4°C. Fatty acids were derivatized using two-step derivatization. First, keto- and aldehyde groups were protected by the reaction with MOX (methoxylamine-HCl in pyridine, 1:2) overnight at room temperature. Then excess derivatizing agent was evaporated and dry residue was converted to MOX-TMS (trimethylsilyl) derivative by reacting with bis(trimethylsilyl) trifluoroacetamide with 10% trimethylchlorosilane (Regisil) at 60°C for 20 min. Resulting MOX-TMS derivatives were analyzed by GC-MS. For the analysis of fatty acid methyl esters, the column temperature was initially set at 100°C and held for one minute, then ramped 8°C/min until 170°C and held for 5 min. Samples were then ramped 5°C/min until 200°C, and held for 5 min. Finally, samples were ramped 10°C/min until 300°C, and held 10 min. Masses were monitored via the SIM acquisition mode. For metabolites, the column temperature was initially set at 80°C and held for one minute, then ramped 5°C/min until 220°C and held for 5 min. Samples were then ramped 5°C/min until 200°C, and held for 5 min. Finally, samples were ramped 10°C/min until 300°C, and held 10 min. Masses were monitored via the SIM acquisition mode. Metabolomics data were analyzed using the MSD ChemStation Software, version: F.01.03.2357 (Agilent Technologies). Metabolite counts were normalized using gamma-hydroxybutyrate.

Untargeted metabolomics was performed using LC-MS. Samples were homogenized in chilled 70% methanol/20% water/10% chloroform. 10 µL of each homogenate was used for protein concentration measurements. The rest was vortexed for 15 seconds and kept on ice for 5 minutes, repeated twice. The homogenates were then centrifuged at 1000x*g* for 15 min at 4° C. The supernatants were dried and re-suspended in 98% water/acetonitrile containing internal standards. Three-microliter aliquots taken from each sample were pooled and the QC standard was analyzed every 6^th^ injection. In addition, we collected MS2 level data on representative control and treated samples. Untargeted metabolomics was performed by injecting 3 µL of each sample onto a 10 cm C18 column (Thermo Fisher, CA) coupled to a Vanquish UHPLC running at 0.3mL/min using water and 0.1% formic acid as solvent A and acetonitrile and 0.1% formic acid as solvent B. The 15-min gradient used is given below. The Orbitrap Q Exactive HF was operated in positive and negative electrospray ionization modes in different LC-MS runs over a mass range of 56-850 Da using full MS at 120,000 resolution. The DDA acquisition (DDA) included MS full scans at a resolution of 120,000 and HCD MS/MS scans taken on the top 10 most abundant ions at a resolution of 30,000 with dynamic exclusion of 40.0 seconds and the apex trigger set at 2.0 to 4.0 seconds. The resolution of the MS2 scans were taken at a stepped NCE energy of 20.0, 30.0 and 45.0. An in-house data preprocessing tool was employed for spectral feature extraction and deconvolution, which includes putative metabolite identification assignment using the National Institute of Standards and Technology Mass Spectral Library (NIST SRD 1A version 17). The spectral features were log-transformed and further analyzed via MetaboLyzer (1) using 0.7 for ion presence threshold, p-value threshold of 0.05 using non-parametric Mann-Whitney U-test, and false discovery rate (FDR) correction set at 0.1 in the positive ESI and at 0.2 in the negative ESI mode. The resulting peak table was further analyzed via MetaboLyzer. First the data was normalized to protein concentration in each sample. The relative abundance values for each spectral feature were then calculated with respect to a labeled internal standard (betaine-d9). The ion presence threshold was then set at 0.7 in each study group for the downstream analysis via MetaboLyzer. Data were then log-transformed, Gaussian normalized, and analyzed for statistical significance via the non-parametric Mann-Whitney U test (FDR corrected p-value <0.1 in positive and <0.2 in negative ESI modes). Ions present in just a subset of samples were analyzed as categorical variables for presence status via the Fisher’s exact test. All p-values were corrected via the Benjamini-Hochberg step-up procedure for false discovery rate (FDR) correction. The data were then utilized for PCA, putative identification assignment, and pathway enrichment analysis via KEGG. In this dataset, 9,322 spectral features were detected, from which 1,422 features were putatively assigned an identification in HMDB within a pre-defined 7 ppm m/z error window. Also, the MS/MS spectra of 304 of these features matched unique compounds in NIST Mass Spectral Library (v17) with a cosine similarity threshold of 0.7.

### In vivo studies

All experiments involving mice were approved by the Case Western Reserve University Institutional Animal Care Regulations and Use Committee (2018-0063). Mice were maintained under pathogen-free conditions in the animal facility with standard chow and nutrient-free bedding. Six-to-eight-week-old, female, athymic nude mice (Foxn1 nu/nu) were purchased from Harlan Laboratories (no. 6903M). For the indicated experiments, hyperglycemia was induced either by allowing mice to consume D30 (dextrose 30% water) or by the administration of streptozotocin (Thermo Scientific, S0130) starting two weeks before cancer cell implantation. The blood glucose levels in streptozotocin-treated mice were titrated to a non-toxic range with daily subcutaneously injections of long-acting insulin glargine (Fisher Scientific, NC0767732). Patient derived xenograft samples were purchased from The Jackson Laboratory (#TM01212) and propagated in nude mice. For flank xenograft experiments, 1×10^6^ cells were suspended in 200µL of a PBS:matrigel solution (1:1) and injected subcutaneously into the right flank. For all flank xenograft experiments, tumor volumes were measured twice per week using a caliper (volume = length x width^2^/2). Body weights were measured weekly. For orthotopic experiments, 4×10^4^ Luciferase-expressing KPC K8484 cells were suspended in 30 µL of a PBS:matrigel solution (1:1) and injected into the pancreas of C57BL/6J mice at 10 weeks of age. Equal numbers of male and female mice were used. Briefly, a 0.5 cm left subcostal incision was made, the tail of the pancreas was externalized, the mixture was carefully injected into the pancreas, and then returned to the peritoneal cavity. On postoperative day 7, the presence of pancreatic tumors was confirmed with bioluminescence imaging after injection of 100 µL intraperitoneal Luciferin (50 mg/mL). Mice with confirmed tumors were then randomized to the indicated treatment conditions. Chemotherapies: gemcitabine (75 mg/kg, twice weekly), FOLFIRINOX (FFX; oxaliplatin 5 mg/kg, 5FU 25 mg/kg, and irinotecan 50 mg/kg, once weekly). For rescue studies, BSO (4.4 g/L water, ad libitum) and NAC (1.2 g/L water, ad libitum) were used.

### Clinical outcome analyses

We retrospectively identified patients who presented with metastatic PDAC at University Hospitals Cleveland Medical Center (2010-2020) and stratified them according to the usage of chemotherapy (vs. supportive care). We utilized raw glucose values extracted from electronic medical records to determine glycemic status for both cohorts. Glucose values were analyzed across two-time intervals: pre-diagnosis (obtained within the 365 days preceding PDAC diagnosis) and during the treatment period (based on chemotherapy initiation date). We stratified patients who received chemotherapy by glycemic status during the treatment period into two groups: high glucose (at least one glucose value ≥200 mg/dL after the initiation of chemotherapy) and normal glucose (all glucose values <200 mg/dL after the initiation of chemotherapy). Identical thresholds were used for patients who did not receive chemotherapy. Both pre- and post-diagnosis treatment values were utilized for stratification of the supportive care cohort due to the limited number of glucose values available. These stratification parameters are in line with American Diabetes Association criteria for diagnosis of diabetes based on random glucose levels (*29*).

## Data Availability

Open source software were used for RNA-seq analysis: FastQC (https://www.bioinformatics.babraham.ac.uk/projects/fastqc/), Trim Galore (http://www.bioinformatics.babraham.ac.uk/projects/trim_galore/), R (v 3.6.3 and v 3.4.2), STAR (v 2.7.0e), DESeq2 (v 1.26.0), and RSEM (v 1.3.2). RNA sequencing data was deposited into Gene Expression Omnibus with accession number GSE194369. All other data supporting the findings of this study are available from the corresponding author on reasonable request.

## Statistical analyses

In vitro data are provided as mean ± s.d. or mean ± s.e.m. from three or more than three independent experiments, respectively. Survival distributions were estimated using Kaplan-Meier estimation and compared by log-rank tests. Pairwise comparison of tumor growth trajectories employed longitudinal mixed models with random intercept and with time viewed as categorical. Box-Cox transformations (log or square root) were used if supported by residual and normal probability plots. For multiple comparisons adjustment, the Holm method was adopted (StataSE v16.0 (Statacorp LLC, College Station, TX) was used for clinical analyses). For demographic and clinical data comparisons between patients in the high and normal glucose groups, continuous variables were compared using the Wilcoxon rank-sum test and categorical variables using Pearson’s chi-squared test. The Nelson-Aalen estimate was used to graphically depict cumulative hazard of death over time. Multivariable Cox proportional hazards regression was used to identify factors associated with overall survival, defined as the time from diagnosis to death or last follow-up. Variables included in multivariable models were those deemed clinically relevant. A p<0.05 was used to indicate statistical significance.

## Acknowledgments

Grant support for this research comes from Postdoctoral Fellowship Grant NCI 1F32CA247466-01 (A.V.- G.), NIDDK R33DK070291, NCI R01CA196643 (H.B.), NCI R37CA237421, R01CA248160, R01CA244931 (C.A.L), UMCCC Core Grant P30CA046592 (C.A.L), NCI P30 CA010815-53, NCI 5 R37 CA227865-04 (J.M.S.), American Cancer Society MRSG-14-019-01-CDD (J.M.W.), 134170-MBG-19-174-01-MBG (J.M.W.), NCI R37CA227865-01A1 (J.M.W.), the Case Comprehensive Cancer Center GI SPORE 5P50CA150964-08 (J.M.W.), Case Comprehensive Cancer Center P30 Core Grant P30CA043703 and University Hospitals research start-up package (J.M.W.). We are grateful for additional support from numerous donors to the University Hospitals pancreatic cancer research program, including the John and Peggy Garson Family Research Fund, The Jerome A. and Joy Weinberger Family research fund, Robin Holmes-Novak in memory of Eugene, and Rosi and Sabi Behar.

